# Regulation of Decision Threshold by the Locus Coeruleus

**DOI:** 10.1101/2025.09.23.678165

**Authors:** Hongjie Xia, Maxime Maheu, Gary A. Kane, Benjamin B. Scott

## Abstract

A fundamental challenge for decision-making under uncertainty lies in regulating the balance between speed and accuracy. Humans and animals solve this problem by adjusting their decision thresholds — the criterion that determines how much information is required before committing to a choice. While brain regions associated with this process have been identified, the neural circuits able to *directly* alter decision thresholds remain to be identified. Here we investigate the role of the locus coeruleus (LC) norepinephrine (NE) system in controlling the balance between speed and accuracy during decision making. Through cell-type specific chemogenetic manipulations, we discovered that LC-NE activation increased decision thresholds. We further demonstrated that this effect is replicated by systemic administration of the α2 adrenergic receptor (α2 AR) agonist, clonidine. Notably, α2 AR activation drove changes in decision threshold specifically, without reproducing other LC-NE activation effects such as promoting task engagement. Together, these results suggest that LC-NE regulates decision thresholds via activation of downstream α2 ARs.

**Highlights:**

- LC-NE activation increases reaction time and accuracy of perceptual decisions
- Behavioral effects are consistent with an elevation of decision thresholds
- Heightened decision thresholds are replicated by selective activation of α2 ARs
- Activation of LC-NE and α2 ARs differently alter task engagement

## Introduction

A core feature of adaptive behavior is the ability to make decisions under uncertainty. In this situation, humans and other animals face a fundamental challenge: deciding when to terminate deliberation and commit to a choice. Prioritizing speed can yield error-prone decisions, whereas emphasizing accuracy delays outcomes. This balance, known as the speed-accuracy tradeoff, is a cornerstone of human and animal behavior in uncertain and time-sensitive environments ^1,2^.

Conceptually, a balance between speed and accuracy can arise from adjustments in decision threshold, defined as the amount of evidence required before making a choice, with higher thresholds yielding slower and more accurate choices ^3^. Past research on the neural implementation of decision thresholds in primates largely focused on cortical structures, reporting involvement of the premotor and cingulate ^4,5^ as well as the superior colliculus ^6^. However, the circuits that alter the threshold remain unresolved. One candidate is the brainstem locus coeruleus (LC) norepinephrine (NE) system, a circuit known to play an important role in behavioral flexibility ^7-9^. Here, we tested the idea that the LC-NE is involved in controlling the threshold for decisions, effectively shifting the balance of speed and accuracy during decision-making.

The locus coeruleus is a small, bilateral nucleus located in the dorsal pons of the hindbrain and serves as the brain’s principal source of norepinephrine. Despite its small size, the LC sends widespread axonal projections to cortical and subcortical regions ^10^, positioning it as a key modulator of brain-wide activity supporting adaptive decision-making ^11-14^. To date, the involvement of the LC-NE system in regulating decision thresholds has been inferred from either pupillometry or pharmacological manipulations in human ^15,16^. Yet pupillometry is not LC-specific ^17^, and many pharmacological agents broadly influence catecholamine signaling ^18^. Consequently, direct evidence for the role of LC-NE in regulating the decision threshold is still lacking.

In this study, we directly investigated how LC-NE system shapes the balance between speed and accuracy during decision-making under uncertainty. We used chemogenetic manipulations of the LC-NE system in rats performing a behavioral task designed to offer a tradeoff between speed and accuracy. LC activation resulted in slower and more accurate decisions, consistent with elevated decision thresholds. These effects were replicated by administration of a selective α2 adrenergic receptors (α2 ARs) agonist, clonidine. Together, these findings suggest that LC-NE activation promotes more deliberative decision-making strategies, likely via α2 AR signaling.

## Results

### Rats trade between speed and accuracy in a free-response perceptual task

We trained rats in a perceptual task designed to elicit a tradeoff between speed and accuracy ^19^. In this task, reward location is indicated by a sequence of light flashes distributed between the left and right nose ports, with the rewarded side having a higher probability of flashes (75% congruent; Fig. 1A). Animals are allowed to respond at any time during the presentation of the flash sequence by poking into either the left or right nose port. Because each flash provides uncertain evidence about the rewarded side, rats can increase their accuracy by sampling more flashes.

**Figure 1.**
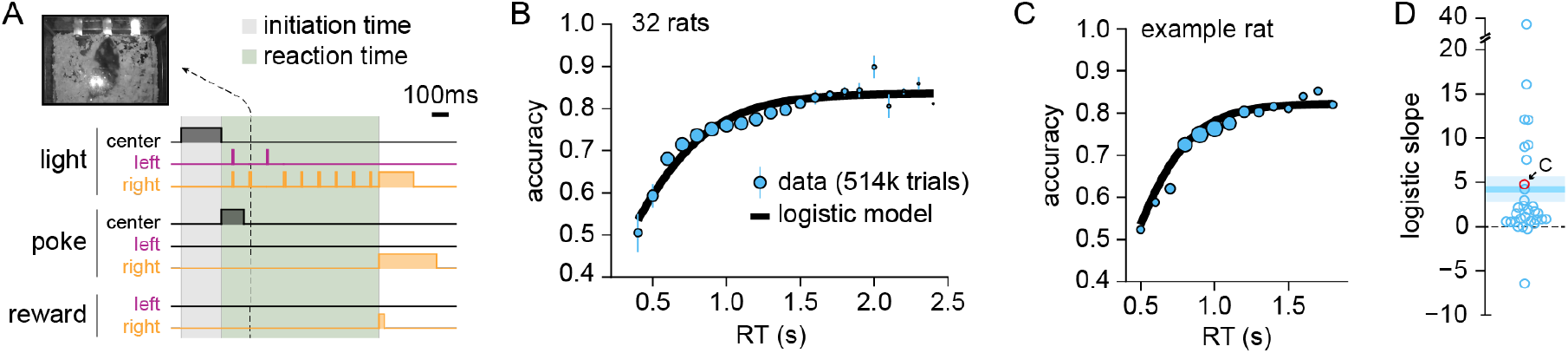
Rats trade between speed and accuracy in a free-response perceptual task. **(A)** Schematic of the task. Each trial begins when the rat nose-pokes into the center port. A sequence of brief flashes (10ms; 10Hz) is then presented from either the left or right ports (after an initial bilateral flash). Each flash has a 75% probability of being presented on the rewarded side. Rats can make a choice at any time by nose-poking into one of the two side ports; point at which flash presentation will stop. Importantly, reward delivery is determined based on the side with the greater generative flash probability, and not based on the actual sequence of flashes, so that rats benefit from sampling from longer durations. ITI: inter-trial interval. **(B)** Chronometric function reporting accuracy as a function of reaction times (RT; 100ms bins; blue dots) for rats (*n* = 32; mean ± SEM). Dot size is indexed on the proportion of trials in each RT bin. Black line shows the prediction of a logistic regression model fitted to rat behavior. **(C)** Accuracy across trials with different reaction times (RT) from an example rat. Data is shown as blue dots, regression fit as black line. **(D)** Slope parameter from the logistic regression model shown in panels B and C. Positive parameter value indicates that accuracy increases with RT. Each dot corresponds to one rat. Rat shown in panel C is indicated with a red circled dot.

Rats performed this task with a group-level accuracy of 75.8% (*n* = 32; range = 68.4-82.8%, s.d. = 0.7%) and median RT of 0.95s (range = 0.72-1.14s, s.d. = 0.14s). Importantly, most rats showed a positive relationship between accuracy and RT, such that waiting longer improved performance (Fig. 1B-D; Fig. S1). These data suggest that rats exhibit a speed– accuracy trade-off in this task, in which longer RTs allow the integration of evidence from multiple flashes.

### Chemogenetic activation of LC-NE shifts the speed-accuracy tradeoff

We tested whether the LC-NE system regulates rats’ speed-accuracy tradeoff in this task using a chemogenetic approach. We injected viruses carrying excitatory DREADDs (hM3Dq) under a synthetic promoter selective to NE neurons (PRS×8 ^20^) into the LC bilaterally (see *Methods*).

We first validated the approach by confirming selective transduction of TH^+^ neurons in the LC (Fig. 2A-B) and increased expression of the immediate-early gene cFos, a proxy for neural activation. Injection of deschloroclozapine (DCZ) ^21^ caused a significant increase in cFos levels in rats with excitatory DREADDs in comparison to a separate group of rats expressing a control transgene (PRS×8-mCherry; Fig. 2C). This data are consistent with previous studies reporting increased LC spiking activity following PRSx8-hM3Dq stimulation in rats ^22^.

**Figure 2.**
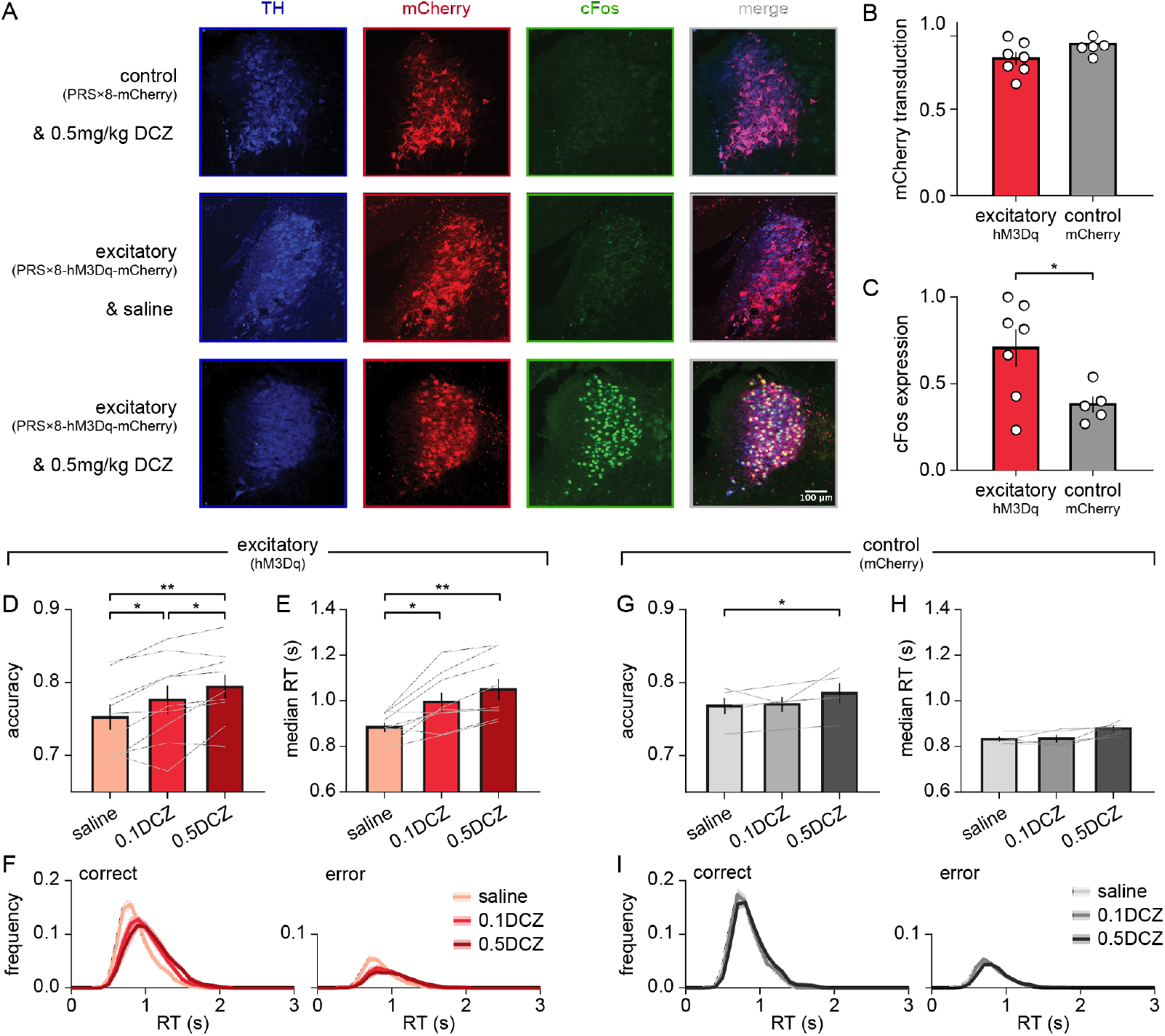
Chemogenetic activation of LC-NE increases accuracy and reaction time. **(A)** Immunofluorescence images of the LC. Top row: control group (PRS×8-mCherry) following injection of 0.5mg/kg DCZ. Middle row: excitatory DREADD group (PRS×8-hM3Dq-mCherry) following saline injection. Bottom row: excitatory DREADD group following 0.5mg/kg DCZ injection. TH: tyrosine hydroxylase. **(B)** Proportion of TH^+^ neurons expressing the transgene (mCherry). **(C)** Proportion of TH^+^ and mCherry^+^ neurons expressing cFos after DCZ injection. Unpaired *t*-test, *t*_(10)_ = 1.155, *p* = 0.035. **(D-F)** Behavioral effects in the excitatory DREADDs group (PRS×8-hM3Dq-mCherry, *n* = 9). **(D)** Mean accuracy (± SEM). Each line represents one animal. Paired *t*-tests with Bonferroni correction, saline vs. 0.1 DCZ: *t*_(8)_ = 3.088, *p* = 0.045; saline vs. 0.5 DCZ: *t*_(8)_ = 4.461, *p* = 0.006. * *p* < 0.05,*\*\* p* < 0.01. **(E)** Median reaction time (mean ± SEM) across animals. Each line represents one animal. Paired *t*-test with Bonferroni correction were performed for saline vs. 0.1DCZ: *t*_(8)_ = 3.431, *p* = 0.027; saline vs. 0.5DCZ: *t*_(8)_ = 4.316, *p* = 0.008; and 0.1DCZ vs. 0.5DCZ: *t*_(8)_ = 3.922, *p* = 0.013. * *p* < 0.05, *\*\* p* < 0.01 **(F)** RT distributions in correct (left) and error (right) trials. Mean ± SEM. Bin width = 0.1 s. **(G-I)** Control group, PRS×8-mCherry (*n* = 5). **(G)** Mean accuracy (± SEM). Each line represents one animal. One way ANOVA, *F*_(2,8)_ = 4.939, *p* = 0.040. Paired *t*-test with Bonferroni correction, saline vs. 0.5DCZ, *t*_(4)_ = 5.774, *p* = 0.013. \**p* < 0.05 for pairwise comparison. **(H)** Median reaction time (mean ± SEM). Each line represents one animal. One way ANOVA, *F*_(2,8)_ = 5.607, *p* = 0.030. **(I)** RT distributions in correct (left) and error (right) trials. Mean ± SEM. Bin width = 0.1 s.

Behaviorally, chemogenetic activation of LC-NE neurons simultaneously slowed RT and improved accuracy in the task in comparison to control saline injections (Fig. 2D-F, Fig. S2). These effects were observed at 0.1mg/kg DCZ and further increased after 0.5mg/kg DCZ administration (Fig. 2D-F, Fig. S2).

By contrast, no changes in accuracy or RT were observed in the group of rats expressing the control transgene (PRS×8-mCherry) after administration of 0.1mg/kg DCZ (Fig. 2G-I). However, we noted that at 0.5mg/kg DCZ, there was a modest change in RT and accuracy (Fig. 2G-H), suggesting that high doses of DCZ may produce a marginal behavioral effect independent of hM3Dq activation. Nevertheless, these effects were smaller than the effects of stimulation with hM3Dq (Fig. S2).

In a separate experiment, DCZ-induced stimulation of inhibitory DREADDs (PRSx8-hM4Di-HA) produced no detectable effect of at either 0.1mg/kg or 0.5mg/kg (Fig. S3).

Together, these data indicate that activation of LC-NE neurons promote slower and more accurate decisions, reflecting a shift in the tradeoff between speed and accuracy.

### Activation of LC-NE increases boundary separation

To directly determine if LC-NE activation alters decision threshold, we accounted for rat behavior using a drift diffusion model (DDM; Fig. 3A). The DDM conceptualizes decision-making as a stochastic process in which a decision variable drifts towards one of two decision boundaries under influence of the stimulus ^23-25^. The DDM provides a unified account of both accuracy and RT and, as such, is a widely adopted framework for describing shifts in the speed-accuracy tradeoff ^26^.

**Figure 3.**
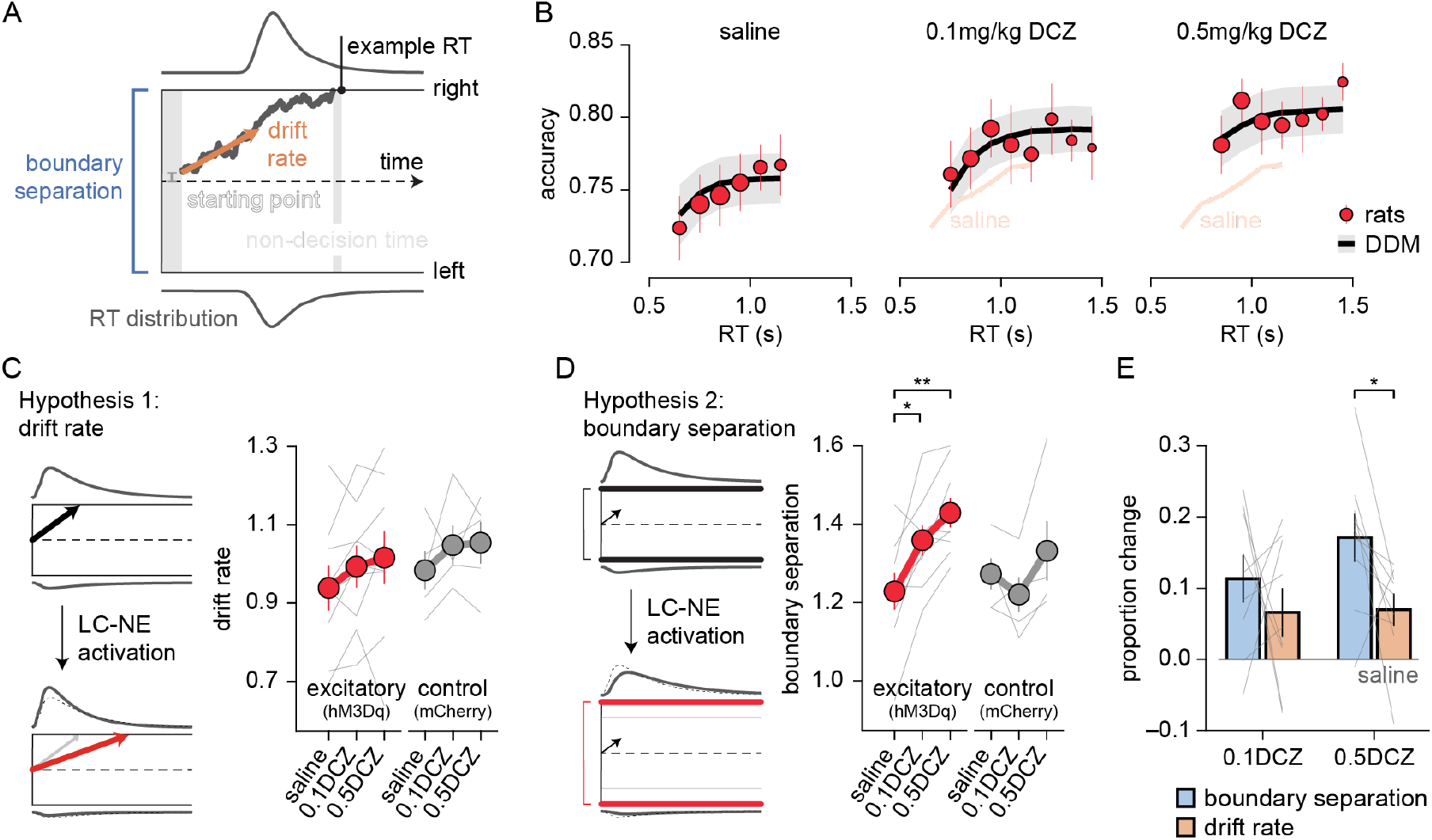
Chemogenetic activation of LC-NE increases boundary separation. **(A)** Schematic of the drift diffusion model (DDM) and its four free parameters: drift rate, boundary separation, starting point bias, non-decision time. **(B)** DDM fit of behavioral data from the excitatory group. Red dots indicate rat accuracy (mean ± SEM) within each RT bin. Black lines show fitted DDM predictions (mean ± SEM). **(C)** Illustration of potential changes in drift rate within the DDM (left). Mean fitted drift rate (± SEM) for the excitatory and control groups (right). **(D)** Illustration of potential changes in boundary separation within the DDM (left). Mean fitted boundary separation (± SEM) for the excitatory and control groups (right). Paired *t*-tests with Bonferroni correction, saline vs. 0.1mg/kg DCZ: *t*_(8)_ = 3.587, *p* = 0.021, and saline vs. 0.5mg/kg DCZ: *t*_(8)_ = 5.598, *p* = 0.002. * *p* < 0.05, ** *p* < 0.01. **(E)** Proportional change in boundary separation and drift rate relative to saline following administration of 0.1mg/kg or 0.5mg/kg DCZ. Paired *t*-tests, saline vs. 0.5mg/kg DCZ: *t*_(8)_ = 2.314, *p* = 0.049.

Free parameters in the DDM include the drift rate, which captures the rate at which the sensory evidence influences the decision variable; and boundary separation, which sets the distance between the two decision thresholds ^27^ (for left and right decisions; see *Methods*) (Fig. S4).

We fitted the DDM separately to data from different experimental conditions and inspected possible changes in model parameters. The DDM captured the speed-accuracy tradeoff in this task well (Fig. 3B). DCZ injections in excitatory DREADDs-expressing rats produced a significant increase in boundary separation in comparison to saline injections; an effect that was DCZ dose-dependent (Fig. 3D, 3E). In contrast, the drift rate parameter remained mostly unaffected by DCZ injections (Fig. 3C). DCZ injections in the group of control (mCherry) rats did not result in significant changes in any of the DDM parameters (Fig. 3C-D, Fig. S5).

Next we considered the alternative hypothesis that the effect of LC-NE activation may reflect a shift in attention, not a change in decision threshold ^28^. To test this hypothesis, we fitted rats’ decisions with a dual-state model which considers that choices are either informed by the flash sequence or made randomly (see *Methods*). Experimental manipulations did not alter the proportion of stimulus-independent decisions, which remained around 5% (Fig. S6).

Together, these data are consistent with a model in which LC-NE neurons regulate decision thresholds, rather than processes peripheral to the decision itself, such as the sensitivity to sensory evidence (as captured by DDM’s drift rate) or attentional states (as captured by a dual-state model).

### Activation of α2 ARs increases accuracy, RT and boundary separation

Our previous experiments demonstrated that LC-NE activation causes a shift in rats’ speed-accuracy tradeoff consistent with an increase in decision threshold. However, an open question concerns the mechanisms mediating this shift. LC-NE neurons release NE as well as other factors, which are known to bind to different receptor subtypes ^29^. Amongst these subtypes, α2 adrenergic receptors (α2 ARs) have the highest affinity towards NE ^30^, and exhibit widespread expression in regions associated with decision-making ^31,32^.

To determine the role of α2 ARs in the shift in speed-accuracy tradeoff, we systemically administered the α2 AR agonist, clonidine (5, 10, 20, 50μg/kg; see *Methods*). In comparison to saline injections, clonidine administration showed a dose-dependent increase in accuracy, with a peak effect at intermediate doses (20μg/kg; Fig. 4A). Similar to LC-NE activation, increases in accuracy were coupled with a slowing of RT (Fig. 4B-C, Fig. S7) and DDM fits revealed a selective increase in boundary separation (Fig. 4D).

**Figure 4.**
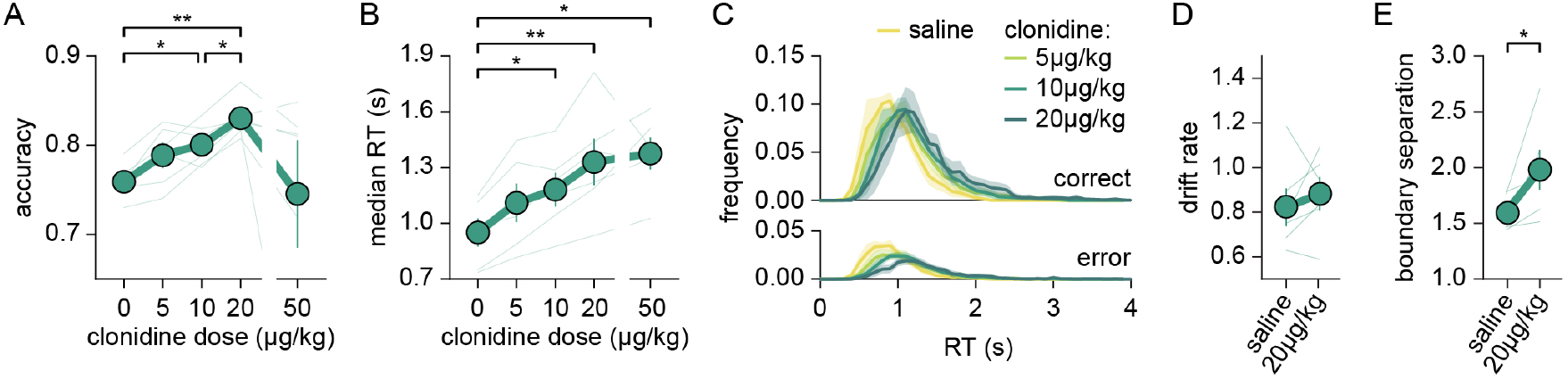
Activation of α2 adrenergic receptors increases boundary separation. **(A)** Mean accuracy (± SEM); *n* = 6. Thin line represents one animal. Paired *t*-test with Bonferroni correction: saline vs. 10μg/kg clonidine, *t*_(5)_ = 6.386, *p* = 0.014; saline vs. 20μg/kg clonidine, *t*_(5)_ = 8.304, *p* = 0.004; 10μg/kg clonidine vs. 20μg/kg clonidine, *t*_(5)_ = 6.095, *p* = 0.017; * *p* < 0.05, ** *p* < 0.01. **(B)** Median reaction time (± SEM). Thin line represents one animal. Paired *t*-test with Bonferroni correction: saline vs. 10μg/kg clonidine, *t*_(5)_ = 7.119, *p* = 0.008; saline vs. 20μg/kg clonidine, *t*_(5)_ = 5.682, *p* = 0.024; saline vs. 50μg/kg clonidine, *t*_(5)_ = 5.616, *p* = 0.025; * *p* < 0.05, ** *p* < 0.01. **(C)** RT distributions (0.1s bins) (± SEM) in correct (top) and error trials (bottom) following administration of saline or clonidine (various doses). **(D)** Mean fitted drift rate (± SEM) and **(E)** mean fitted boundary separation (± SEM) from DDM under saline and 20μg/kg clonidine conditions. Paired *t*-test, *t*_(5)_ = 2.653, *p* = 0.045; * *p* < 0.05.

At the highest doses used (50μg/kg), clonidine impaired performance, including a reduction in the total number of trials completed per session and yielded high inter-individual variability in accuracy (Fig. 4A, Fig. S7). Together these data indicate that moderate activation of α2 ARs mimics the effects of LC-NE activation on shifting the speed-accuracy tradeoff and suggest that the LC may alter decision thresholds via release of NE and activation of α2 ARs.

### Manipulations of LC-NE change task engagement

Past research points towards an involvement of LC-NE in task engagement ^7,33,34^, suggesting its role may extend beyond shifting the speed-accuracy tradeoff and elevating thresholds during decision-making. In this task, engagement can be assessed by initiation time: the latency to initiate a new trial (Fig. 1A).

Compared to saline injections, DCZ-driven LC-NE activation significantly shortened initiation times; an effect that was not observed in the control (mCherry) group (Fig. 5A-B). As trial initiation is typically faster following rewarded trials ^35^, reduced initiation times following LC-NE activation could simply reflect the higher accuracy observed in this condition (Fig. 3). Instead, initiation times were shorter upon LC-NE activation following both correct and error trials, indicating a simultaneous effect of LC-NE stimulation in shifting the speed-accuracy tradeoff and enhancing task engagement (Fig. 5A).

**Figure 5.**
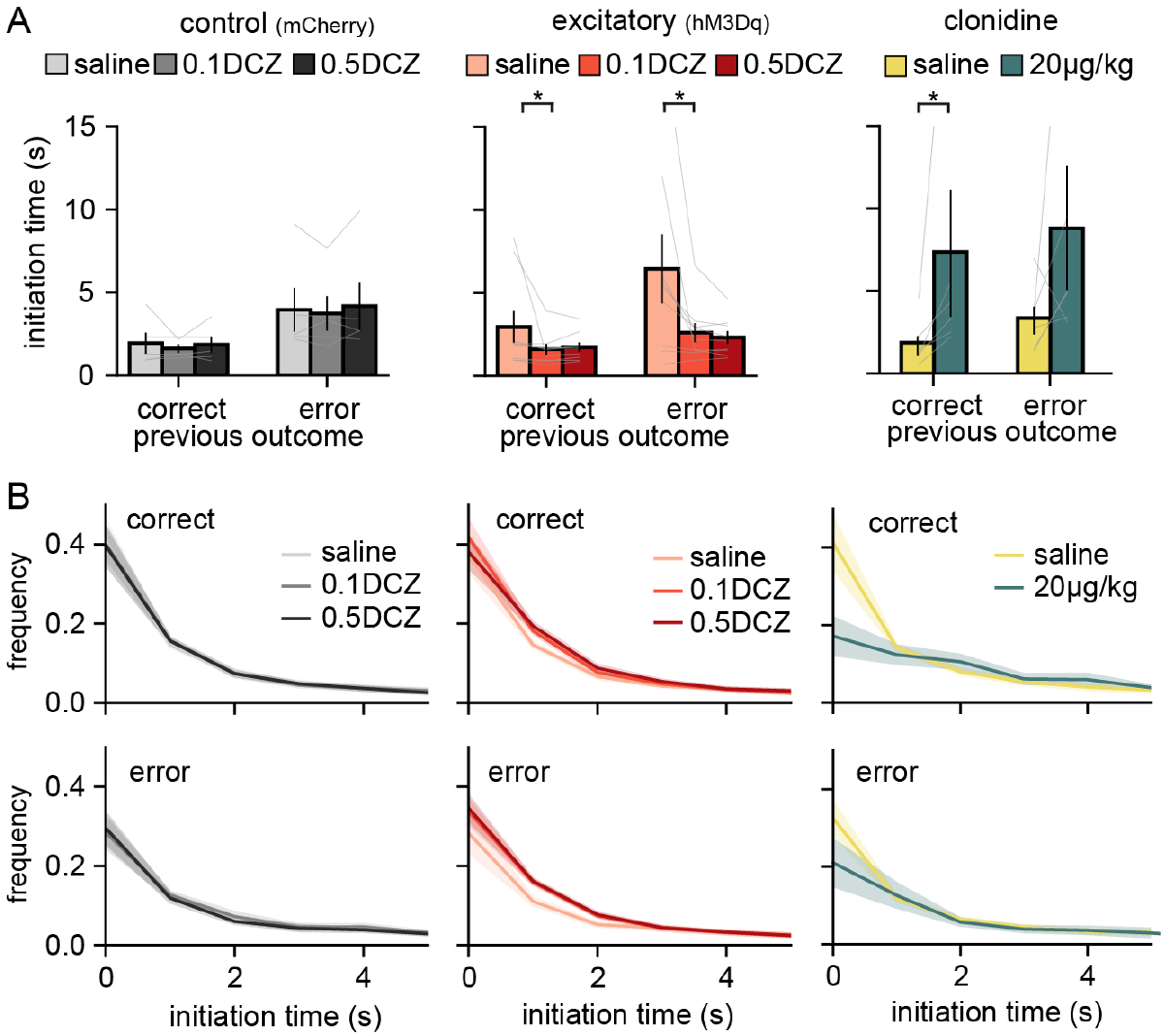
Manipulations of LC-NE change task engagement. **(A)** Median initiation time (Mean ±SEM) by previous trial outcome (correct vs. error). Each line represents one animal. Left: control group, *n* = 5. Middle: excitatory group, *n* = 9. Wilcoxon signed-rank test: saline_correct_ vs. 0.1DCZ_correct_, *p* = 0.023; saline_error_ vs. 0.1DCZ_error_, *p* = 0.035. Right: clonidine group, *n* = 6. Wilcoxon signed-rank test: saline vs. 20μg/kg clonidine, *p* = 0.031. * *p* < 0.05. **(B)** Initiation time distributions (bin size = 1s; mean ± SEM) following correct (top) or error (bottom) trials in control (left), excitatory (middle), and clonidine (right) groups. The distribution aligns to left edge of each bin, such that at 0 represents the frequency from 0-1s.

Thus, activation of LC-NE neurons induced a non-trivial pattern of results: slower RTs during the stimulus period but faster initiation latency at trial onset. This suggests that slower RTs after LC-NE activation is not the mere consequence of unspecific motor slowing. Instead, these observations point to a role for the LC-NE system in flexible adjustments of the deliberative process including the elevation of decision thresholds.

We observed a different pattern of results following clonidine administration. By comparison to saline injections, administration of clonidine slowed initiation times (Fig. 5A-B); an effect which scaled with dose (Fig. S7). Thus, activation of α2 ARs had the opposite effect on trial initiation compared with direct LC-NE activation. These data suggest that LC-NE enhancement of engagement may rely on circuit mechanisms involving receptor types beyond α2 ARs.

## Discussion

In this study, we investigated the role of LC-NE neurons in setting the balance between speed and accuracy during decision-making under uncertainty. We found that chemogenetic activation of LC-NE neurons drives an increase in decision thresholds and that this change could be reproduced by pharmacological activation of α2 ARs.

These results suggest a new role for the LC-NE in decision making. Previous models of LC-NE function have emphasized its role in a range of cognitive processes, including attention ^13,36^, working memory ^37-39^, and behavioral flexibility ^9,33,40-42^. During perceptual decision making, LC-NE has also been linked to modulation of sensory gain ^13,43-45^; an effect that is typically captured by changes in the drift rate within the DDM ^36^. In this study, we observed no significant change in drift rate but instead observed a change in boundary separation, which reflects the decision threshold. Our free-response task emphasized speed–accuracy tradeoff rather than fine sensory discrimination^14,36^, and the use of highly salient stimuli in our task may have reduced our ability to detect LC-NE-dependent changes in sensory gain. Another possibility is that LC-NE may differentially regulate distinct aspects of perception and decision making depending on the spatio-temporal pattern of neuromodulator release^7,46^. For instance, decision thresholds may be influenced by tonic LC activity, whereas phasic activity may preferentially support the regulation of sensory processes^13,44,47^.

Prevailing theory emphasizes LC heterogeneity^10,48-50^ including cell-type specific ^51,52^ or projection-specific populations ^53,54^, which may differentially influence cognition. Recent findings suggest how specific projections of LC neurons may play a role in the decision threshold. In particular, the superior colliculus (SC), a region implicated in the control of decision thresholds ^6^ is innervated by the LC ^55,56^ and abundantly expresses α2 ARs ^31,57^. Direct suppression of SC neurons, by NE binding to α2 ARs could delay burst activity that reflects decision commitment, Alternatively, α2 ARs could influence decision thresholds by acting in regions upstream of the SC such as the cortex ^56^ or the substantia nigra (SN) ^58^. When cortical drive is weak, SN inhibition dominates, suppressing SC activity and delaying commitment. Because adrenergic receptors are enriched in long-range pyramidal neurons of frontal cortex ^59^, activation of α2 ARs could reduce excitatory output to the SC, thereby increasing decision thresholds. In both scenarios, LC-NE signaling may gate the timing of decision commitment via its influence on SC excitability.

While we demonstrate a role of α2 ARs in the regulation of decision thresholds, we did not directly investigate the influence of other receptors subtypes. Thus is possible that additional receptor subtypes (including those specific to factors others than NE) might be involved in supporting the effects observed here. Indeed, β AR blockade has been reported to alter the amount of evidence gathered prior to choice ^16^, suggesting that adrenergic mechanisms beyond α2 ARs could influence decision thresholds. Together, these studies raise the possibilities that multiple receptor subtypes and forebrain circuits may jointly regulate decision threshold.

In addition to increasing decision threshold, we also found that LC-NE is involved in task engagement. Other receptors may play a role in mediating increased engagement. Such receptors could include β and α1 ARs, and possibly dopamine receptors, given that LC-NE has been reported to co-release dopamine ^29,60-62^. These findings are consistent with the idea that the behavioral consequences of LC-NE involve distinct contributions from multiple receptor subtypes and co-transmitters.

## Conclusions

By combining chemogenetic, pharmacological manipulation and behavioral modeling, we discovered that direct activation of the LC-NE system increases decision thresholds; an effect likely mediated through NE release and binding of α2 ARs. We also found that direct LC-NE activation promotes task engagement, whereas α2 ARs produced the opposite effect. Together, this work provides a clear behavioral link between LC-NE signaling and decision threshold, offering a new insight into how LC-NE shapes cognition and complex behavior.

## Acknowledgement

We thank Christa Rose for help with daily rat training. This work was supported by NIMH award number R56MH132732 to BBS, a Marie Skłodowska-Curie postdoctoral fellowship to MM, and a Center for Systems Neuroscience distinguished fellowship award to GAK.

## Author contributions

HX, GAK and BBS designed the experiments. HX collected the data. HX and MM analyzed the data. HX and MM prepared the figures. HX wrote the manuscript. BBS, HX and MM edited the manuscript.

## Methods

### Animals

All experiments and procedures were performed in accordance with protocols approved by the *Boston University Animal Care and Use Committee*. Adult male and female Long-Evans rats (*Charles River Laboratory*; criver.com) aged 3-6 months were used. Rats were food restricted and maintained above 80% of baseline weight throughout training period. Rats were housed in 12hr light-dark cycle. All behavioral training were conducted during light period.

### Behavioral task

Rats performed the task in customized operant chamber for two hours daily, from Monday to Friday. Each chamber contains three nose ports equipped with a white LED. This task has been previously described ^19,63^. To briefly summarize here, rats begin each trial by poking their nose in the center port. This triggers a first bilateral flash followed by a 10Hz sequence of 10ms unilateral flashes, occurring from either the left or the right port. Occurrence of flashes followed a Bernoulli process with a 75% probability of a flash occurring on the correct side, and a 25% probability on the incorrect side. The correct and incorrect side is randomly selected on each trial. Rats report their choice by poking to either side port at any time during the trial. Following correct trials, rats receive 25μl of a 10% sucrose solution and are imposed a 5s inter-trial interval. Following incorrect trials, rats receive a 3s timeout punishment, leading to a total of 8s inter-trial interval. If rats fail to respond within 8s following trial initiation, the trial is considered an “omission”. In practice, omitted trials were rare (1.09%). A total of 32 rats (16 females) performed the task without chemogenetic or pharmacological manipulations (Fig. 1), yielding a dataset of 514,978 trials.

### Behavioral training

In the first stage of training, rats are rewarded by poking the illuminated side port. In stage 2, rats are rewarded by first poking into the center port, then to the illuminated side port. In stage 3, all flashes appear on the correct side and rats received a reward after poking on that side. In stage 4, the probability of flashes occurring on correct vs. incorrect is set to 90%-10%. In stage 5, this probability is set to 80%-20%. Stage 6 is the final stage and correspond to a probability of 75%-25%. All trials in this study were collected in stage 6.

### Surgery

A total of 24 rats (13 females) were included in the chemogenetic experiment. Once rats reached the final stage of training, they received a bilateral virus injection in the LCs. Viruses used were either adeno-associated virus (AAV) obtained from the *UPenn Viral Core* or canine adenoviruses (CAVs) obtained from the *Plateforme de Vectorologie de Montpellier* (<u>plateau-igmm.pvm.cnrs.fr</u>). The surgical procedure started by anesthetizing the rat with isoflurane (5% induction, 1-2% maintenance). For analgesia, buprenorphine (0.02 mg/kg, intraperitoneal) was administered. Rats were aligned in the stereotaxic frame such that lambda was 2 mm above bregma (corresponding to 15-20° head angle). Bilateral burr holes were drilled at the following coordinates: AP (from lambda): −3.9mm, ML: ±1.35, viruses were injected at depth 6-6.5mm from the surface of the brain. Rats received AAV9-PRS×8-mCherry (control), CAV-PRS×8-hM3Dq-mCherry (excitatory group, 5.7×10^12^ pp/mL before dilution, 1:6 diluted with sterile PBS), or CAV-PRS×8-hM4Di-HA (inhibitory group, 12.5×10^12^ pp/mL before dilution, 1:15 diluted with sterile PBS). A total volume of 1μl was injected into each LC at a rate of 100nL/min using a Hamilton syringe. Rats were given 7 days to recover from surgery (before behavioral sessions resume) during which they were given *ad libitum* access to food.

### DCZ administration

DREADDs were activated by deschloroclozapine dihydrochloride (DCZ), which was chosen for its high selectivity for DREADDs ^21^. We obtained DCZ through *NIMH Chemical Synthesis and Drug Supply Program*. DCZ was diluted in sterile saline with a stock concentration of 0.5mg/mL. The stock solutions were kept at −20°C. DCZ stock solution were further diluted to 0.1mg/mL in order to obtain the desired 0.1mg/kg dosage. Working concentrations were kept at room temperature for up to 4 days. To habituate rats to injections, we injected rats with saline during the first week of training. Then, rats were administered with either 0.1mg/kg DCZ, 0.5mg/kg DCZ or saline (i.p.) on interleaved days, 10 minutes prior to the start of the behavioral session. A total of at least five sessions for each concentration were collected from each rat.

### Histology

Rats were euthanized with sodium pentobarbital euthanasia solution at a dosage of, or greater than, 200mg/kg. Rats were perfused with 10% neutral buffered formalin one and a half hour after the start of final training session. Brains were extracted and postfixed with 10% formalin for at least 72 hours before sectioning. Brains were sliced at 40µm thickness using a vibratome. 1 in 3-series sections were incubated overnight with primary antibodies diluted in 2% non-fat milk and 0.1% Triton X-100 in 1xPBS. To confirm the expression of hM4di-HA, a rabbit anti-HA antibody (*Cell Signaling*, C29F4, 1:500) were co-immunostained with a mouse anti-TH antibody (*Immunostar*, 22941, 1:1000). To confirm the increased cell activity following DCZ injections in hM3Dq-expressing animals, brain slices were co-immunostained with a rabbit anti-cFos antibody (*Abcam*, ab190289, 1:500) and a mouse anti-TH antibody. On the next day, after three washes of 1xPBS, slices were incubated with secondary antibodies at room temperature for two hours with agitations. Goat anti-mouse 647 (*Abcam*, ab150115), goat anti-rabbit 488, (Invitrogen, A-11008), and goat anti-mouse 568 (*Invitrogen*, A-11004) secondary antibodies were used during this process. Finally, slices were washed three times with 1xPBS and mounted onto microscope slides.

### Image acquisition and cell counting

All confocal images were taken using a *Nikon C2 Si* confocal microscope. Z-stacks were acquired and channel-specific maximum projection images were obtained using *ImageJ*. Images were then converted into 8-bit gray scale and imported into *CellPose 2*.*0* for segmentations ^64^ (github.com/MouseLand/cellpose). For each channel, 50% of the images were manually labeled and a new model was trained using these manually defined cell masks. This model was then applied to the other half the images. Missing or false positive cell masks were manually curated. Cell masks were opened with *ImageJ*’s ROI manager for counting cells. In order to count overlapped cell masks in any two channels (TH^+^ and cFos^+^ on one hand, and TH^+^ and mCherry^+^ on the other hand), cell masks from each channels were first ‘Combine(OR)’ into a single cell mask and overlapped cell maks were created between two channels using the ‘AND’ function. The total number of cell masks and corresponding fluorescence values were exported for further analyses using custom-made scripts and cell masks smaller than five pixels were eliminated.

### Clonidine administration

A total of 6 female rats were included in this experiment. Clonidine hydrochloride was first dissolved in sterile saline at 1mg/mL concentration, then aliquoted and stored at −20°C (*Tocris*, cat no. 0690). Clonidine was further diluted into a final injection volume of 1mL/kg weekly before usage and stored at room temperature. Rats received intraperitoneal injections of either saline or clonidine (5, 10, 20, or 50μg/kg) on interleaved days. Injections occurred 10 minutes prior to the start of the behavioral session.

### Trial exclusion

Prior to all behavioral analyses, we excluded trials with reaction times (RT) shorter than 0.3s, indicative of false nose port entries (due to photogate instability). In addition, we excluded the 10% of the trials associated with longest RTs on an individual basis; with the idea that behavior in these trials did not reflect stimulus-driven decisions, but often corresponded to unintended initiation.

### Logistic regression model

A logistic regression model explaining accuracy as a function if RT is defined as:

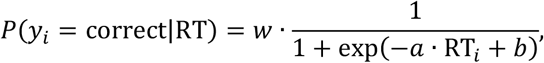

where *y*_*i*_ is the outcome at trial *i*. Free parameters (*w, a, b*) are fitted using the *sklearn* package in *Python* and the slope parameter (*a*) was tested against zero to test for the existence of a speed-accuracy tradeoff in rats

### Drift diffusion model

We accounted for rats’ behavior using a vanilla drift diffusion model (DDM), which casts decision-making as a stochastic process in which a decision particle drifts over time until a decision threshold is reached. The DDM has 4 free parameters: drift rate (*v*), decision boundary (*a*), non-decision time (*t*_0_) and starting point (*z*) ^24,25^. The DDM considers that the stochastic drift follows a Wiener diffusion process:

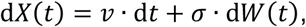

where *X*(*t*) is the decision variable at time *t* and σ is the noise scaling factor. In this model, a decision is made when *X*(*t*) reaches one of two boundaries. The lower boundary is set to 0 and the upper boundary is equal to *a*. The starting point *z* is defined as a proportion of the boundary separation, such that an unbiased process corresponds to *z* = 0.5. Reaction time (RT) is composed of decision time (DT) and non-decision time (*t*_0_), with the latter capturing processes peripheral to the decision process, such as sensory encoding and motor execution:

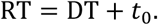

Free parameters are fitted using a custom *R* package (github.com/gkane26/rddm). Parameters are estimated separately for each rat and each condition using the *qmpe* package, which compares observed and simulated RT distributions using a quantile maximum estimation method ^65^. To confirm the recoverability of the DDM, we generated 50 independent sets of parameters by drawing each parameter from a normal distribution, with mean and standard deviation corresponding to the group average from the saline condition With 2000 trials per simulation, similar to those contributed by each rat per condition, the DDM’s free parameters were recoverable (Fig S4).

### Dual-state model

In the dual-stage model, choice outcome is modeled as a mixture of an engaged, logistic process and a stimulus-independent, lapse process. In different versions of the model, choice outcome probability in the engaged state is a function of either stimulus strength (balance between number of left vs. right flashes; see Fig.S1) or RT. In all cases, the probability of a correct choice on trial *i* is given by:

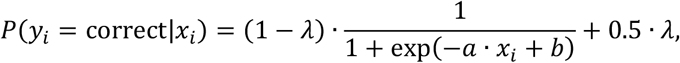

where *x*_*i*_ is the predictive variable (stimulus strength or RT), *a* is the slope and *b* is the intercept of the logistic function. Lapse trials occur with probability *λ* producing random choices independent of *x*_*i*_.

### Statistical analysis

ANOVAs and paired *t*-test were performed using the *pingouin* package in *Python*. To control for family-wise type 1 error, Bonferroni correction was used to adjusted *p*-values. Normality of paired differences was assessed using the Shapiro-Wilk test. For initiation time, most conditions significantly deviated from normality (*p* < 0.05); therefore, we used non-parametric Wilcoxon signed-rank significance tests.

## Supplementary figures

**Supplementary Figure 1.**
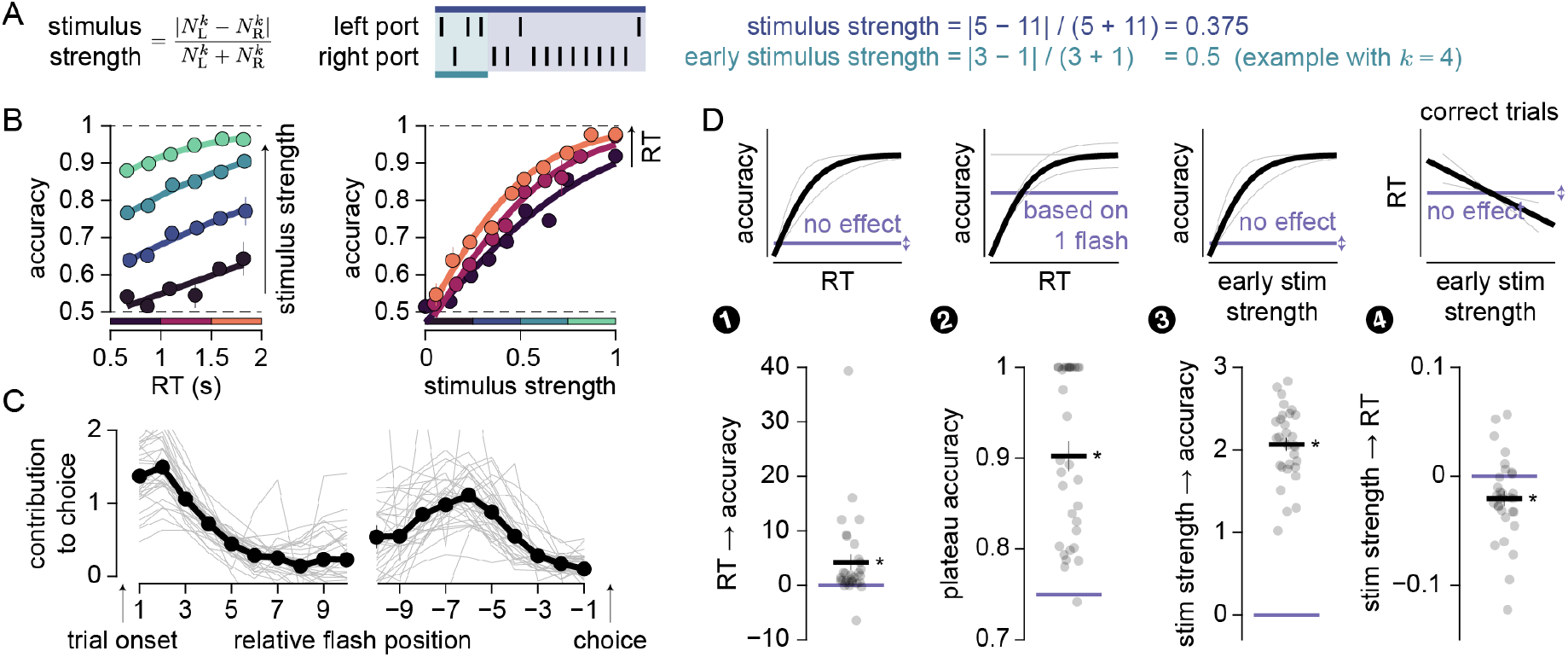
Behavioral signatures of sensory evidence accumulation in rats performing the task. **(A)** The stochastic nature of stimulus presentation yields trials with different stimulus strengths, quantified by the proportion of flashes pointing towards the same location (irrespective of left or right). Stimulus strength can be computed on all the flashes (dark blue) or restricted to the first *k* flashes (light blue; here with *k* = 4) to reflect early commitment to choice. **(B)** Mean ± SEM choice accuracy reflects the combined influence of stimulus strength (blue color coded) and reaction time (RT; red color coded). Thick lines correspond to logistic regression fits and bin limits are shown as colored squares at the bottom of each plot **(C)** The relative contribution (A.U.) of individual flashes (left or right) delivered at different position relative to trial onset (left) or choice (right) is estimated using a logistic regression. Thin individual lines correspond to individual rats. **(D)** Coefficients from logistic (1-3) and linear (4) regression models (depicted in top row) indicate that accuracy increases with RT (1; vs. 0: *t*_(30)_ = 2.97, *p* = 0.006; same as Fig. 1D), plateau accuracy is larger than expected from a single flash strategy (2; vs. 75%: *t*_(30)_ = 9.25, *p* < 0.001), accuracy increases with early stimulus strength (3; *k* = 4; vs. 0: *t*_(30)_ = 26.4, *p* < 0.001), and RT accelerates with early stimulus strength (4; *k* = 4; vs. 0: *t*_(30)_ = −3.00, *p* = 0.005). * *p* < 0.05.

**Supplementary Figure 2.**
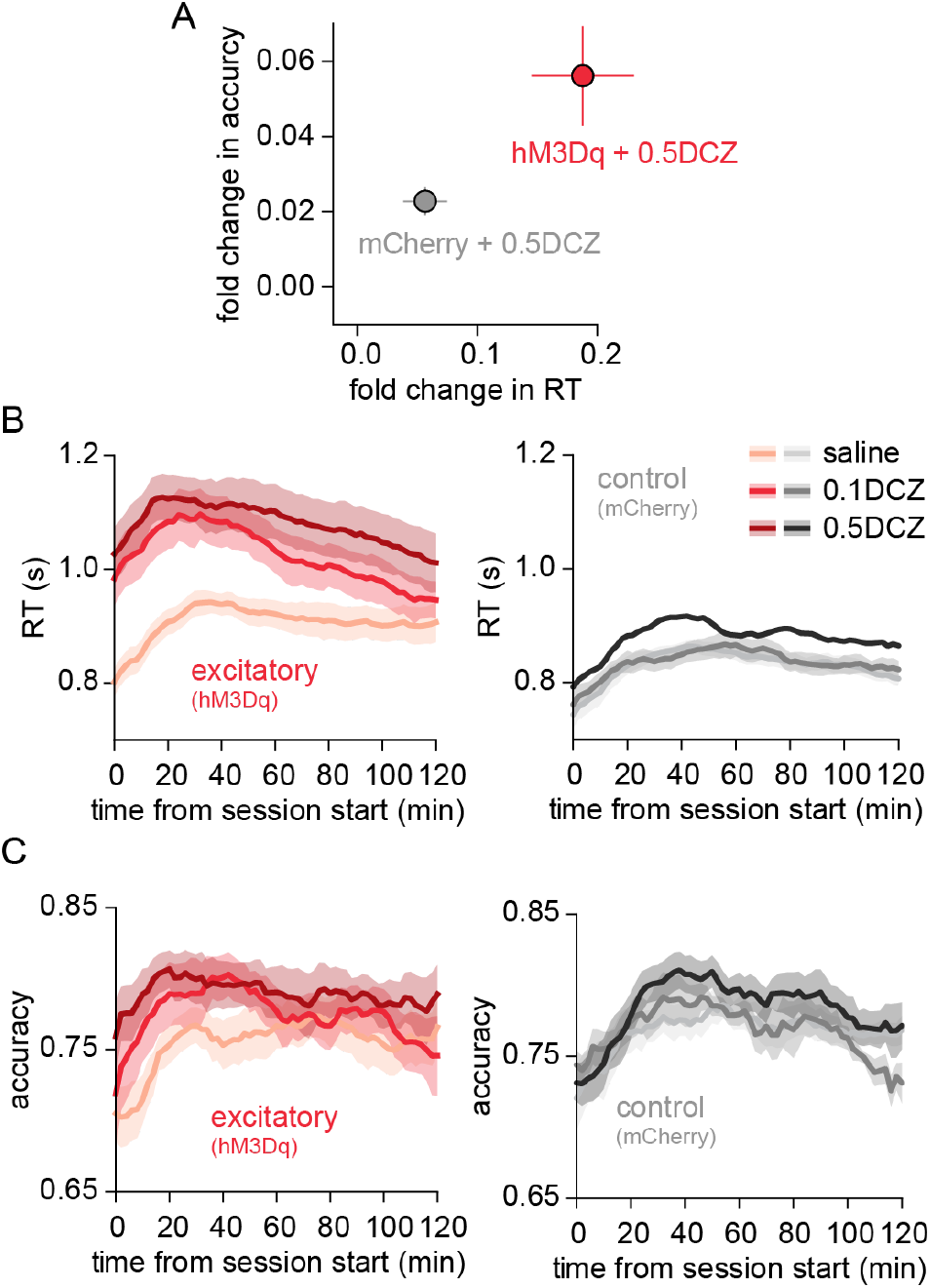
Chemogenetic excitation induced a greater change in behavior than DCZ alone. **(A)** Proportional change of RT and accuracy from saline condition in 0.5mg/kg DCZ in excitatory (red) and control groups (gray). **(B)** Rolling average of RT using a 20-minute window across session in excitatory group (left) and control (right). Shaded area represents SEM across animals. **(C)** Rolling average of accuracy using a 20-minute window across session in excitatory group (left) and control (right). Shaded area represents SEM across animals.

**Supplementary Figure 3.**
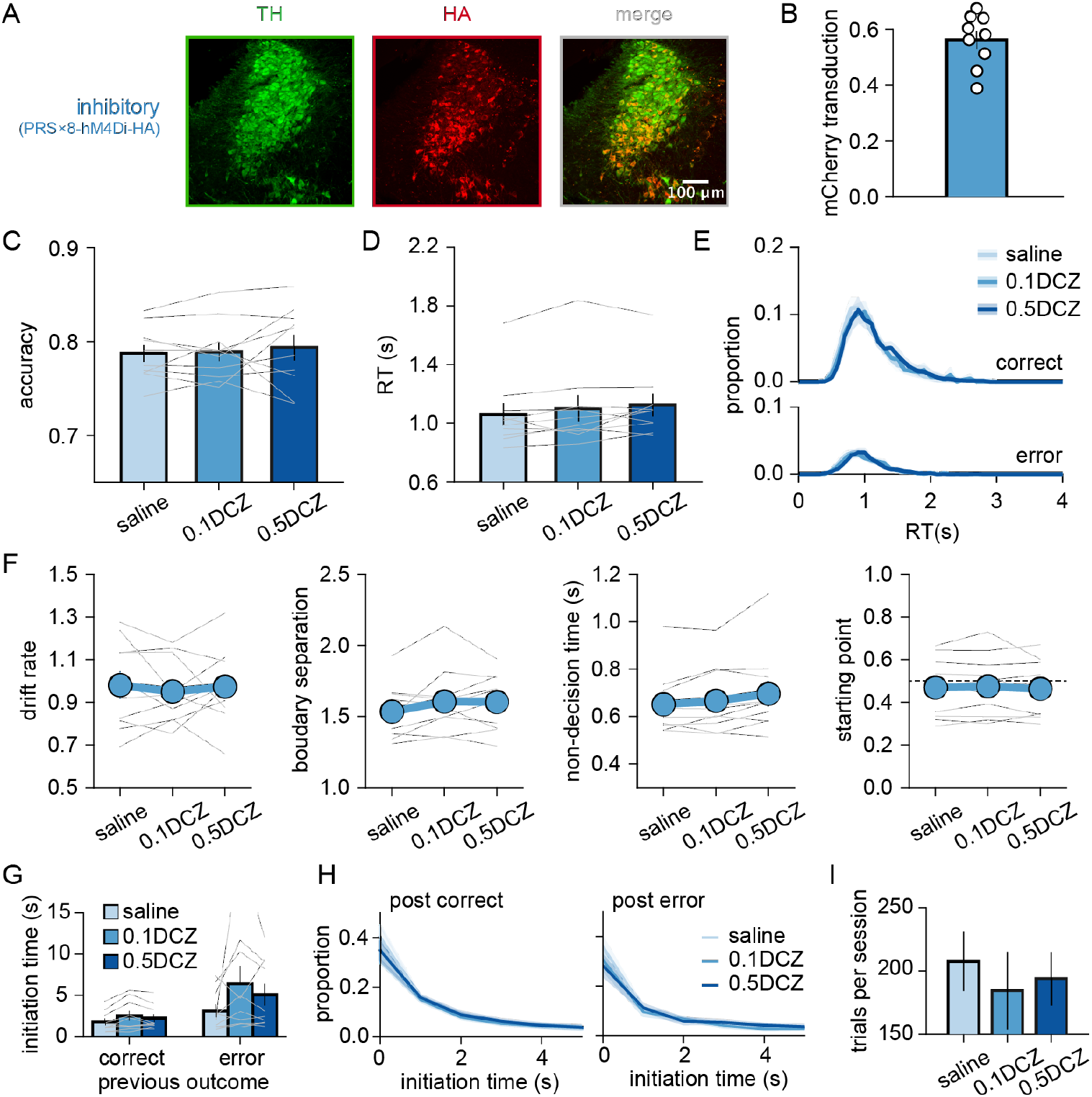
Chemogenetic inhibition of LC-NE does not alter speed and accuracy in the task. **(A)** Immunohistology of rats injected with inhibitory transgene, PRS×8-hM4Di-HA. Green: Tyrosine hydroxylase (TH), Red: HA-tag. **(B)** Mean ± SEM proportion of TH+ neurons expressing the transgene, i.e. HA+. Each circle represents an individual rat. **(C)** Average accuracy across rats (*n* = 10 rats). Thin lines correspond to individual rats. Mean ± SEM. **(D)** Average of median RT across rats. **(E)** RT distributions from chemogenetic inhibitory group. **(F)** Estimated DDM parameter for chemogenetic inhibitory group. **(G)** Average of median initiation time (± SEM) across animals. **(H)** Mean distribution of initiation times sorted by the outcome of previous trials that are correct (left) and error (right) across animals. **(I)** Average number of trials completed per session across animals (± SEM).

**Supplementary Figure 4.**
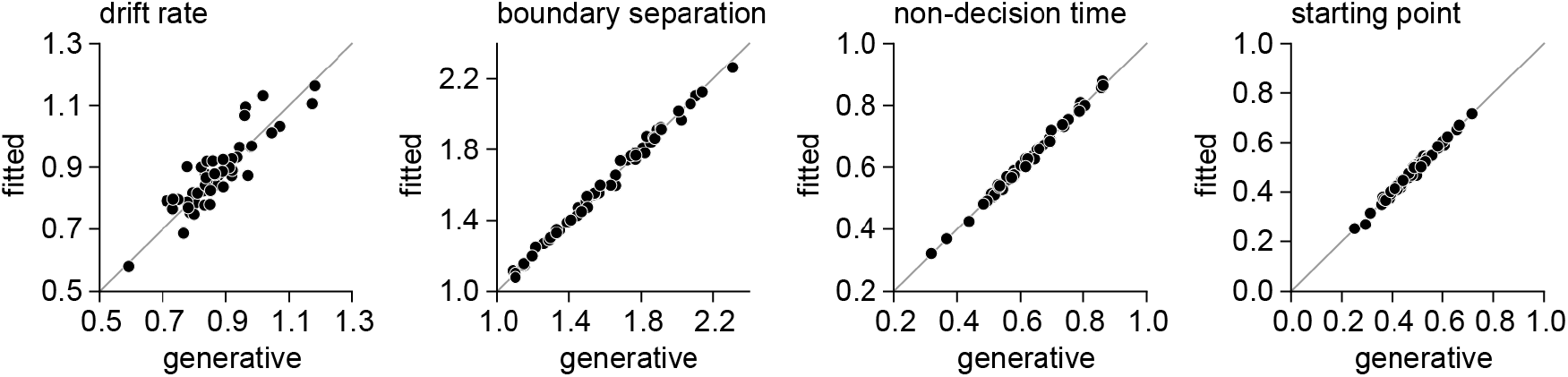
DDM parameter recovery and goodness of fit estimates. Model recovery of generative parameters across 50 artificial agents, each with 2000 trials.

**Supplementary Figure 5.**
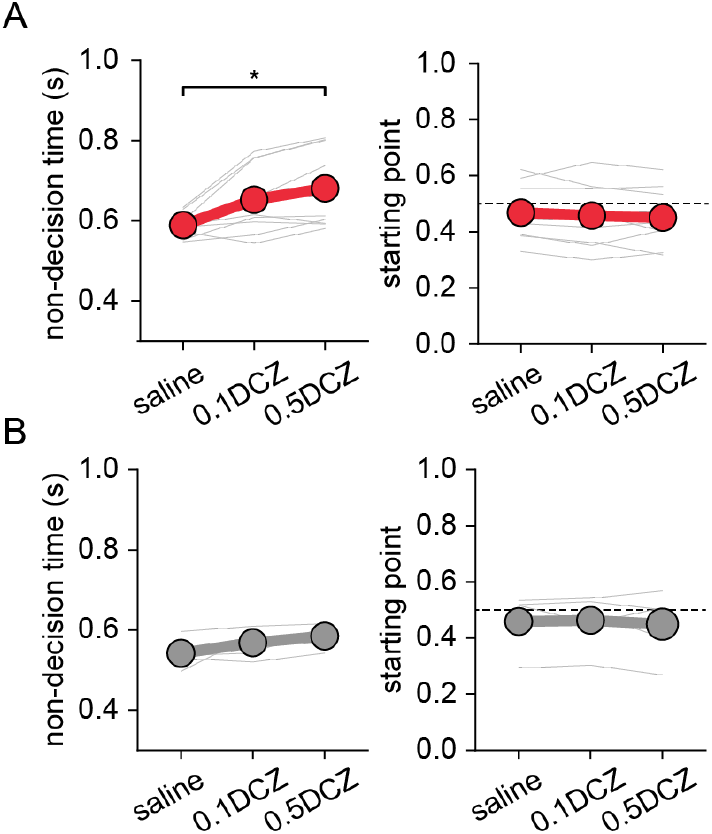
DDM estimates of non-decision time and starting point. **(A)** Excitatory group. Paired t-test with Bonferroni correction, saline vs. 0.5DCZ: *t*_(8)_ = 3.265, p = 0.034. * *p* < 0.05 **(B)** Control group.

**Supplementary Figure 6.**
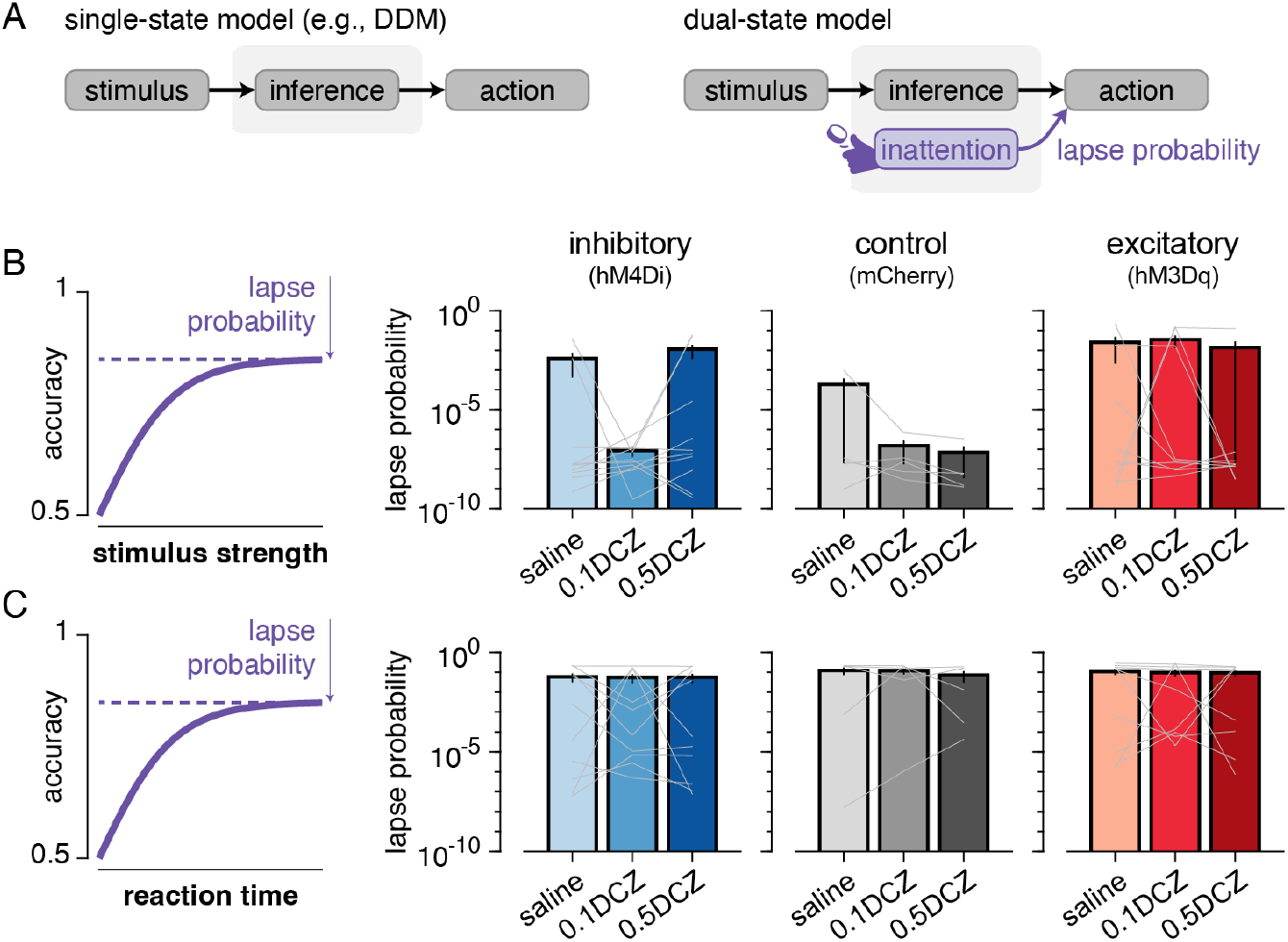
LC-NE manipulations did not detectibly shift attentional allocation. **(A)** Schematics depicting a single-state model in which all choices are stimulus-informed (left) vs. a dual-state model in which choices are either stimulus-informed or made randomly (right). In the dual-state model, the frequency with which random choices are made is controlled by lapse probability parameter. **(B)** Lapse probability is estimated based on a version of the dual-state model relating accuracy to stimulus strength (One-way ANOVA, all *p* > 0.18). **(C)** Same as panel B but from a version of the dual-state model relating accuracy to reaction times (One-way ANOVA, all *p* > 0.59).

**Supplementary Figure 7.**
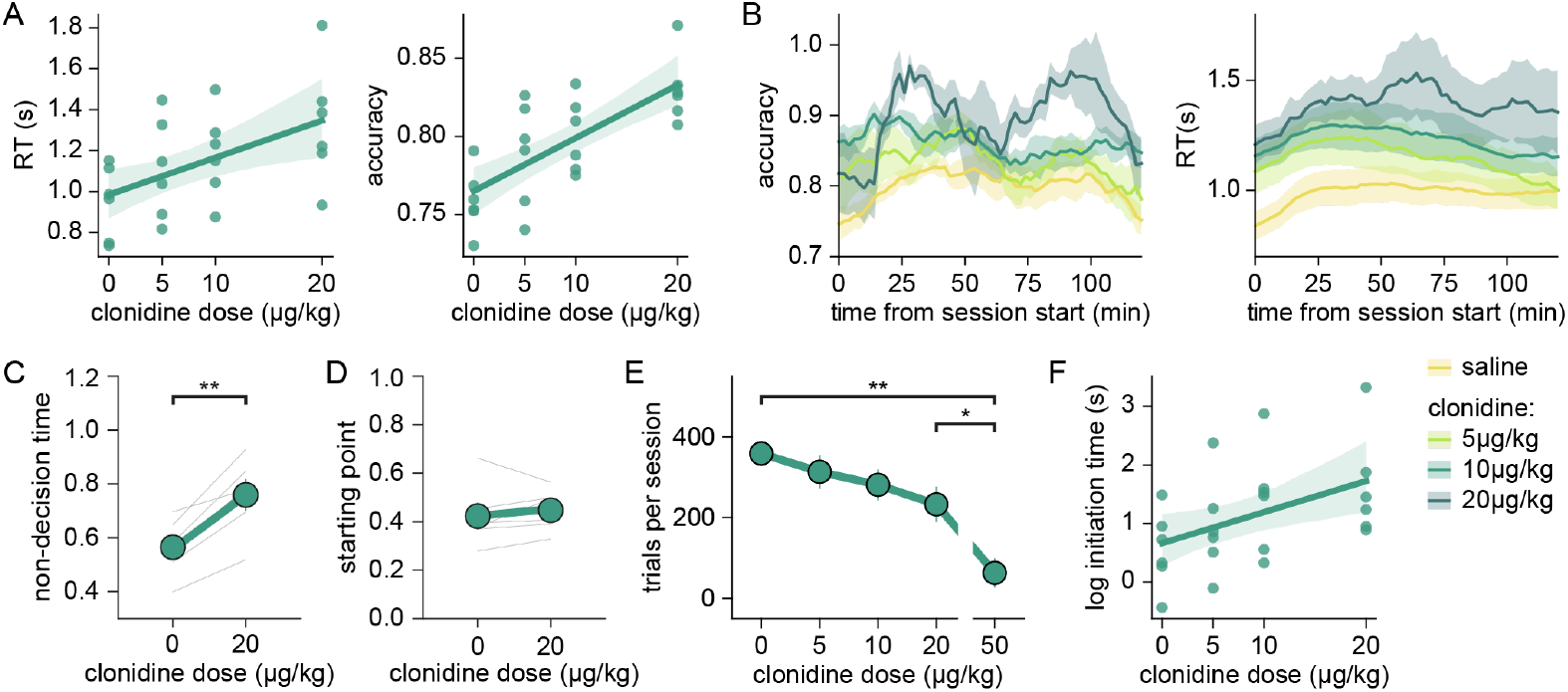
Effect of clonidine, an α2 AR agonist, on the task. **(A-B)** Linear regression model was fitted to **(A)** RT, *R*^2^ = 0.28, *p* = 0.008 and **(B)** accuracy, *R*^2^ = 0.48, *p* < 0.0001, across different clonidine dosages. Each dot represents median RT or median accuracy of a single rat across sessions in the experiment. *n* = 6. **(C)** Rolling average of RT and accuracy using a 20-minute window across sessions. Shaded area represents SEM across animals. **(D)** Estimated non-decision time for saline and 20μg/kg clonidine from DDM. Each line represents individual animal. Paired *t*-test, saline vs. 20μg/kg clonidine: *t*_(5)_ = 5.934, *p* = 0.002, ** *p* < 0.01. **(E)** Estimated starting point for saline and 20μg/kg clonidine from DDM. **(F)** Average number of trials per session across animals. Paired t-test with Bonferroni correction, saline vs. 50μg/kg clonidine: *t*_(5)_ = 8.775, *p* = 0.003. 20μg/kg vs. 50μg/kg clonidine: *t*_(5)_ = 5.251, *p* = 0.033 * *p* < 0.05, ** *p* < 0.05. **(G)** Linear regression fit of log initiation time across rats. Each dot represents individual rats. *R*^2^ = 0.206, *p* = 0.026.

